# Regulation of Liver Regeneration by hepatocyte O-GlcNAcylation in mice

**DOI:** 10.1101/2021.10.14.464401

**Authors:** Dakota R. Robarts, Steven R. McGreal, David S. Umbaugh, Wendena S. Parkes, Manasi Kotulkar, Sarah Abernathy, Norman Lee, Hartmut Jaeschke, Sumedha Gunewardena, Stephen A. Whelan, John A. Hanover, Natasha E. Zachara, Chad Slawson, Udayan Apte

## Abstract

The liver has a unique capacity to regenerate after injury in a highly orchestrated and regulated manner. Here we report that O-GlcNAcylation, an intracellular post-translational modification (PTM) regulated by two enzymes, O-GlcNAc transferase (OGT) and O-GlcNAcase (OGA), is a critical termination signal for liver regeneration (LR) following partial hepatectomy (PHX). We studied liver regeneration after PHX on hepatocyte specific OGT and OGA knockout mice (OGT-KO and OGA-KO), which caused a significant decrease (OGT-KO) and increase (OGA-KO) in hepatic O-GlcNAcylation, respectively. OGA-KO mice had normal regeneration, but the OGT-KO mice exhibited substantial defects in termination of liver regeneration with increased liver injury, sustained cell proliferation resulting in significant hepatomegaly, hepatic dysplasia and appearance of small nodules at 28 days after PHX. This was accompanied by a sustained increase in expression of cyclins along with significant induction in pro-inflammatory and pro-fibrotic gene expression in the OGT-KO livers. RNA-Seq studies revealed inactivation of hepatocyte nuclear 4 alpha (HNF4_α_), the master regulator of hepatic differentiation and a known termination signal, in OGT-KO mice at 28 days after PHX, which was confirmed by both Western blot and IHC analysis. Furthermore, a significant decrease in HNF_α_ target genes was observed in OGT-KO mice, indicating a lack of hepatocyte differentiation following decreased hepatic O-GlcNAcylation. Immunoprecipitation experiments revealed HNF4α is O-GlcNAcylated in normal differentiated hepatocytes. These studies show that O-GlcNAcylation plays a critical role in the termination of LR via regulation of HNF4_α_ in hepatocytes.

**Layman summary:** O-GlcNAcylation is a protein modification that plays a critical role in various biological processes including cell proliferation, differentiation, and disease progression. These studies show that O-GlcNAcylation in hepatocytes is essential for proper liver regeneration. Without O-GlcNAcylation, hepatocytes keep on proliferating eventually forming liver tumors.

## Introduction

The liver has a remarkable regenerative capacity after injury or surgical resection (1). Liver regeneration (LR) is tightly regulated by a plethora of redundant signals that regulate initiation and termination of cell proliferation as well as tissue remodeling. The most widely used model to study liver regeneration (LR) is partial hepatectomy (PHX) where approximately 2/3rd of the liver is surgically removed (2). Liver cells, starting with hepatocytes followed by other cells, enter the cell cycle and undergo a synchronized cell division to restore the lost liver mass. In rodents, this process takes between 3 to 5 days after which proliferation decreases, the newly divided cells undergo redifferentiation and tissue remodeling takes place. These latter events including inhibition of cell proliferation, redifferentiation and tissue remodeling, are termed termination of liver regeneration. Whereas the initiation signals of LR are well characterized, the mechanisms of termination of LR remain understudied (3). Proper termination of liver regeneration is essential because the loss of proper termination signals results in hepatomegaly, defects in redifferentiation of hepatocytes leading to either loss of liver function and liver failure or rapid preneoplastic changes in the liver leading to chronic liver disease.

O-GlcNAcylation is an intracellular post-translational modification (PTM) that involves the addition of a single N-acetylglucosamine (GlcNAc) molecule on an exposed serine or threonine amino acid of a protein. The O-GlcNAcylation cycle is regulated by two enzymes, O-GlcNAc transferase (OGT) and O-GlcNAcase (OGA). OGT catalyzes the transfer of GlcNAc from its carrier molecule UDP-GlcNAc to the protein. Whereas OGA catalyzes the hydrolysis and removal of the GlcNAc motif from the protein (Fig. S1A). In some cases, the residues that undergo O-GlcNAcylation on the target protein can also become phosphorylated providing the cell an additional mechanism of regulating downstream signaling (4). O-GlcNAcylation homeostasis is critical for healthy cells and aberrant O-GlcNAcylation has been linked to various diseases, including cancer, non-alcoholic fatty liver disease (NAFLD) and alcoholic steatohepatitis (ASH) (5-7). O-GlcNAcylation plays a major role in a myriad of cellular processes, including cell proliferation (8-10). Despite the role of O-GlcNAcylation in cell proliferation, little is known about its role in hepatocyte LR. In this study, we investigated the role of O-GlcNAcylation in the regulation of LR after PHX using control, OGA and OGT hepatocyte-specific knock-out mice (OGA-KO and OGT-KO respectively). Our studies revealed that decreasing O-GlcNAcylation leads to impaired termination of LR. Moreover, this is due to the loss of hepatocyte nuclear 4 alpha (HNF4_α_) function, a nuclear receptor critical for hepatocyte differentiation and function.

## Materials and Methods

### Animal Care and Surgeries

All animal studies were approved by and performed in accordance with the Institutional Animal Care and Use Committee at the University of Kansas Medical Center. OGT^fl/Y^ mice were developed by Dr. Natasha Zachara at Johns Hopkins School of Medicine (11). OGA^fl/fl^ mice were developed by Dr. John Hanover at the NIDDK (12). Two-month-old male OGT^fl/Y^ and OGA^fl/fl^ mice were injected intraperitoneal with AAV8-TBG-GFP or AAV8-TBG-CRE (Vector Biolabs) to generate control or hepatocyte specific OGT-KO or OGA-KO animal, as previously stated (13). PHX surgeries were performed on 8-week-old male C57BL/6J, OGT^fl^, OGT-KO, OGA^fl/fl^, and OGA-KO mice as previously described (14). Mice were euthanized at various time points between 0 to 28 days after PHX to collect blood and liver samples. Serum was isolated from clotted blood by centrifugation at 5,000 rcf for 10 minutes at 4°C. A section of the liver was fixed in 10% neutral buffered formalin for 48 h and then paraffin embedded. A piece of liver was then cryopreserved in OCT. Liver injury was measured using serum ALT activity (Pointe Scientific ALT Assay by Fisher Scientific).

### Statistical Analysis

For all experiments not associated with RNA-seq or metabolomics, such as ALT measurements, results are expressed as mean ± standard error of the mean. GraphPad Prism 8 was used to graph and calculated statistics. Student’s t-test or ANOVA with Tukey’s post-hoc was applied to all analyses with a p-value <0.05 being considered significant. Dot plots and heatmaps were produced in RStudio (R version 4.0.3; RStudio Team).

All other methods are described in detail in the supplementary materials.

## Results

### Decrease O-GlcNAcylation resulted in hepatomegaly, liver Injury and defective termination of liver regeneration

Hepatocyte-specific OGT-KO and OGA-KO mice were successfully generated by injecting OGT^fl/Y^ and OGA^fl/fl^ mice with AAV8-TBG-CRE, respectively (Fig 1A-B). OGT^fl/Y^ and OGA^fl/fl^ mice treated with AAV8-TBG-GFP were used as controls (referred to as WT). As expected, deletion of OGT decreased global liver O-GlcNAcylation (Fig. 1A) and deletion of OGA enhanced O-GlcNAcylation levels after PHX (Fig. 1B). Further, deletion of either OGT or OGA decreased the reciprocal enzyme levels (Fig. 1A-B). A significant liver injury as demonstrated by serum ALT levels was observed at 2 days after PHX in WT mice compared to OGT-KO mice. However, OGT-KO mice exhibited significantly higher liver injury at 14 and 28 days after PHX (Fig. 1C). Most importantly, the liver weight to body weight ratio indicated substantial hepatomegaly in the OGT-KO mice at 0, 14 and 28 days after PHX (Fig. 1D). At 28 days after PHX, the liver weight to body weight ratio was 80% higher in OGT-KO mice as compared to its 0-hour level indicating defective termination of regeneration. Interestingly, liver injury or liver weight to body weight ratios were not different between WT and OGA-KO mice indicating that loss of OGA does not affect liver regeneration (Fig. 1E-F).

**Figure 1.**
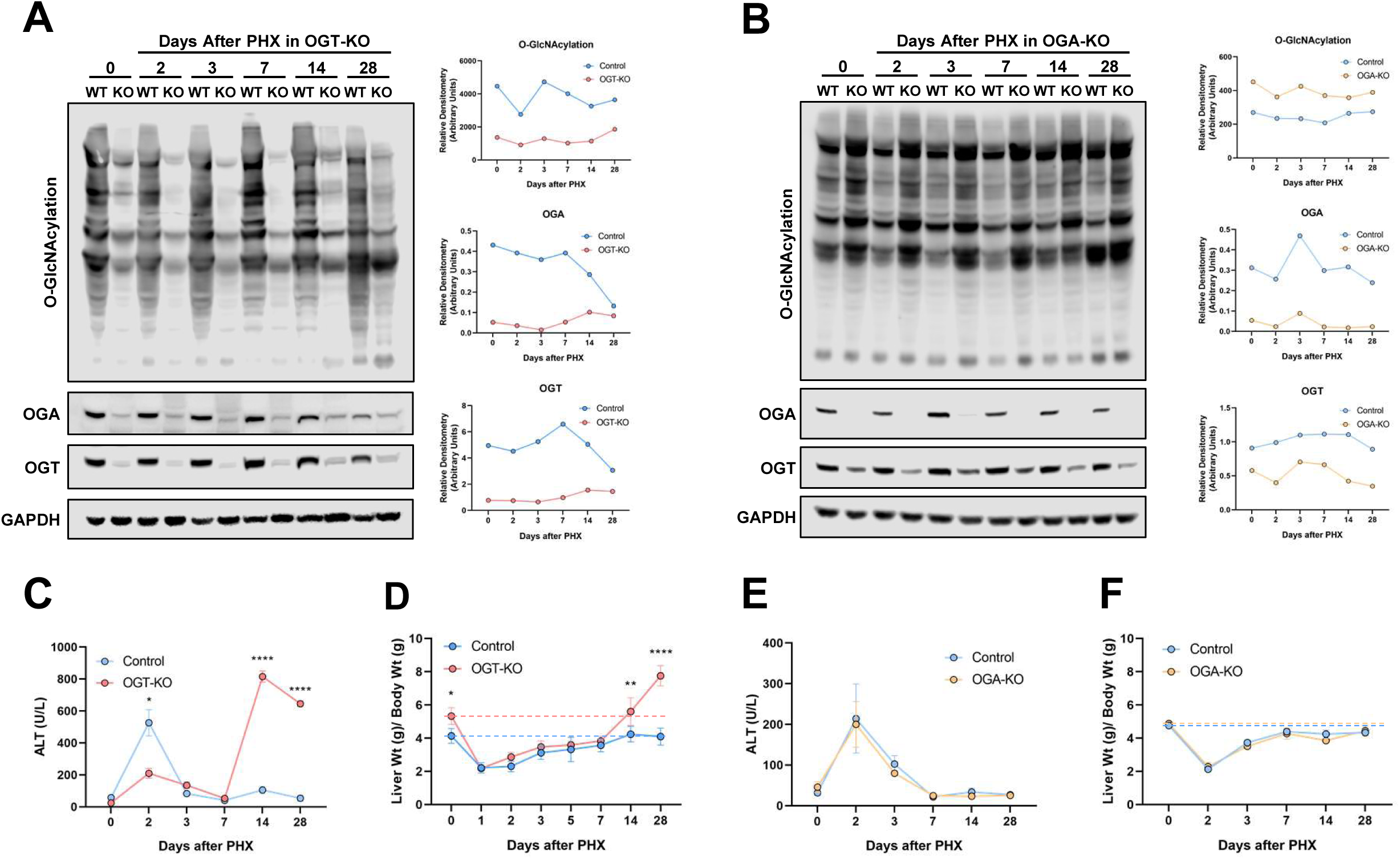
Decreased Hepatocyte O-GlcNAcylation lead to significant increase in liver weight to body weight ratios and liver injury. Western blot analysis of (A) OGT-KO and (B) OGA-KO mice of total liver O-GlcNAcylation levels, OGA, OGT and GAPDH over a time course of 0-28 days after PHX with corresponding densitometry normalized to the loading control. Protein was pooled from 3-5 mice. Line graphs show serum ALT levels in (C) OGT-KO mice and (E) OGA-KO mice and liver weight to body weight ratio for (D) OGT-KO and (F) OGA-KO mice at various time points after PHX. Data are mean ± SEM, * p < 0.05, ** p < 0.01 and **** p < 0.0001.

### Mechanisms of liver injury on OGT-KO mice after PHX

We performed TUNEL staining to determine the mechanisms of cell death in OGT-KO mice at 7, 14 and 28 day time points due to the extent of liver injury. At 14 and 28 days after PHX, the majority of the staining was localized in the cytosol with very limited nuclear staining indicating necrotic cell death and some apoptosis (Fig. S2A). Western blot analysis confirmed activation of both necroptosis and apoptosis at 14 and 28 days after PHX (Fig. S2B-C). Pyroptotic cell death mechanisms did not contribute to cell death determined by western blot analysis (Fig. S2D). Interestingly, OGT-KO mice exhibited elevated p62, an autophagy marker (Fig. S2E). Taken these data together, OGT-KO mice exhibited liver injury due to necroptosis and apoptosis.

### Decreased O-GlcNAcylation resulted in sustained cell proliferation and hepatic dysplasia

We determined cell proliferation in the liver after PHX using PCNA immunohistochemistry. PCNA expression peaked at 2 days after PHX as expected in both WT and OGT-KO (Fig. 2A-B). After day 3, PCNA expression gradually declined in the WT, whereas it remained high in the OGT-KO mice till 28 days after PHX (Fig. 2A-B). Cell proliferation peaked in the WT and OGA-KO mice at 2 days after PHX before declining. Consistent with the liver weight to body weight data, we did not observe any difference in cell proliferation between WT and OGA-KO mice after PHX (Fig. S3A).

**Figure 2.**
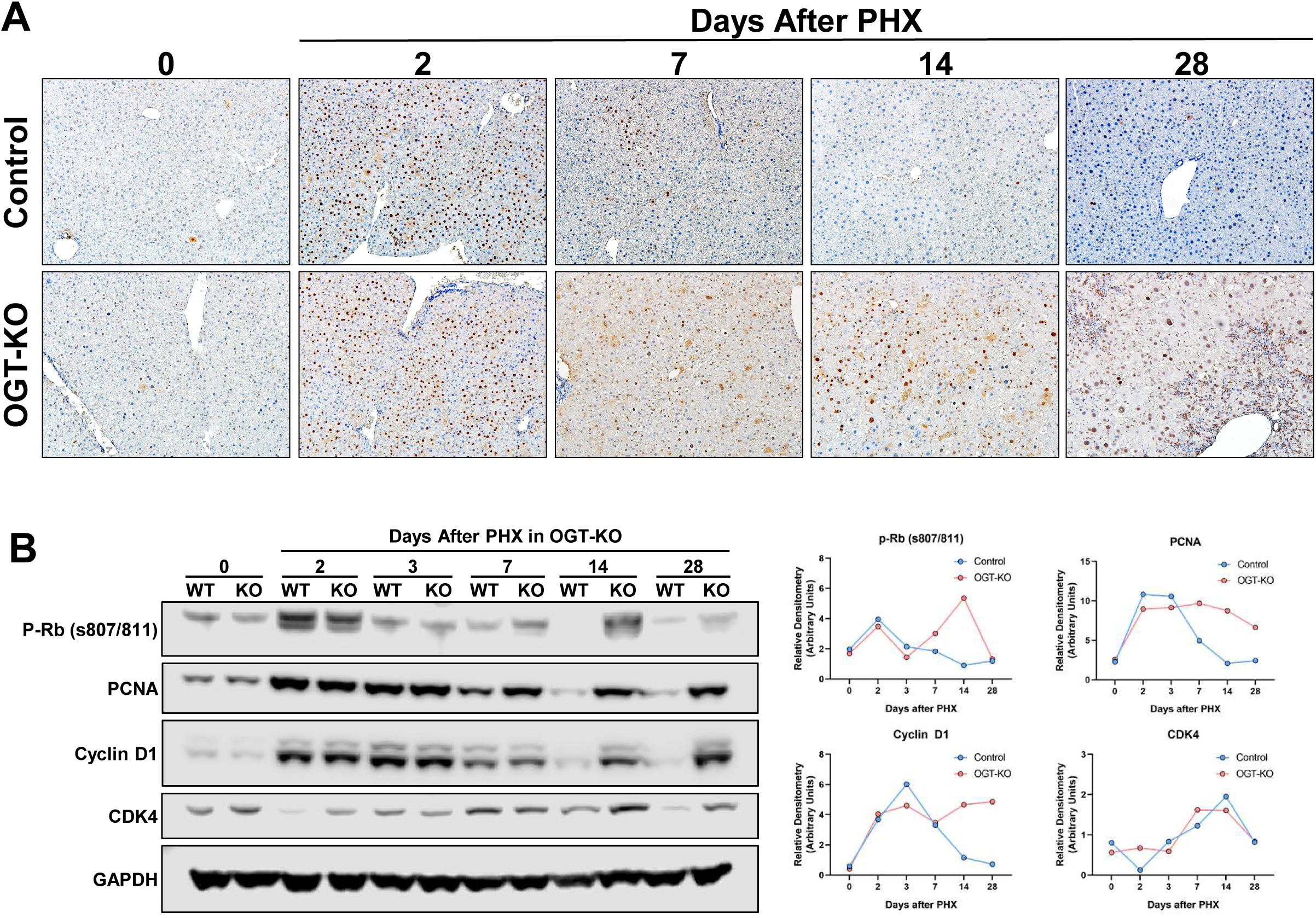
Sustained cell proliferation in OGT-KO mice after PHX. (A) Representative photomicrographs (200x) of PCNA-stained liver sections from OGT-KO and control mice throughout the PHX time course. (B) Western blot analysis of the pro-mitogenic factors p-Rb, PCNA, Cyclin D1 and CDK4 (left panel) in pooled (n=3-5) OGT-KO liver lysates at all time points post-PHX accompanied with the calculated densitometry normalized to GAPDH (right panel).

Next, we investigated the expression of core cell cycle machinery that drives cell proliferation. Western blot analysis revealed that expression of cyclin D1, CDK4 and phosphorylated Rb (p-Rb) protein increased after PHX to similar levels in WT and OGT-KO livers at 2 days after PHX. A steady decline in expression of these proteins was observed in WT mice after the 2-day time point. However, OGT-KO mice exhibited sustained induction in cyclin D1, p-Rb and CDK4, 14 and 28 days after PHX. (Fig. 2B). In OGA-KO mice, expression of these proteins increased at 2 days post-PHX and decreased thereafter. There was no difference between the OGA-KO and WT mice, consistent with cell proliferation data (Fig. S3B).

### Hepatic dysplasia, inflammation and early fibrosis occurred in OGT-KO 28 days after PHX

Hematoxylin and Eosin (H&E) staining was utilized to determine histopathological changes. As expected, OGA-KO mice had little to no effect in histological changes at all time points after PHX (Fig. S3C-D). In contrast, OGT-KO mice manifested significant histological changes at 14 and 28 days after PHX characterized by ballooning hepatocytes (Fig. 3A-B). At 28 days after PHX, the OGT-KO liver showed presence of hepatic nodules containing mitotic figures, surrounded by extensive inflammatory and ductular cells (Fig. 3B). qPCR analysis revealed an induction of Cyclin D1, A2, and B1 gene expression and no change in Cyclin E2 in OGT-KO mice 28 days after PHX (Fig. 3C). Similarly, protein levels of Cyclin D1, A1, E1, and B1 were elevated 28 days after PHX (Fig. 3D). Western blot analysis revealed a significant increase in expression cell proliferation marker PCNA, (Fig. 3E), corroborating. Additionally, PCNA immunohistochemistry (IHC) showed proliferation of both hepatocytes and the ductular cells (Fig. 2C).

**Figure 3.**
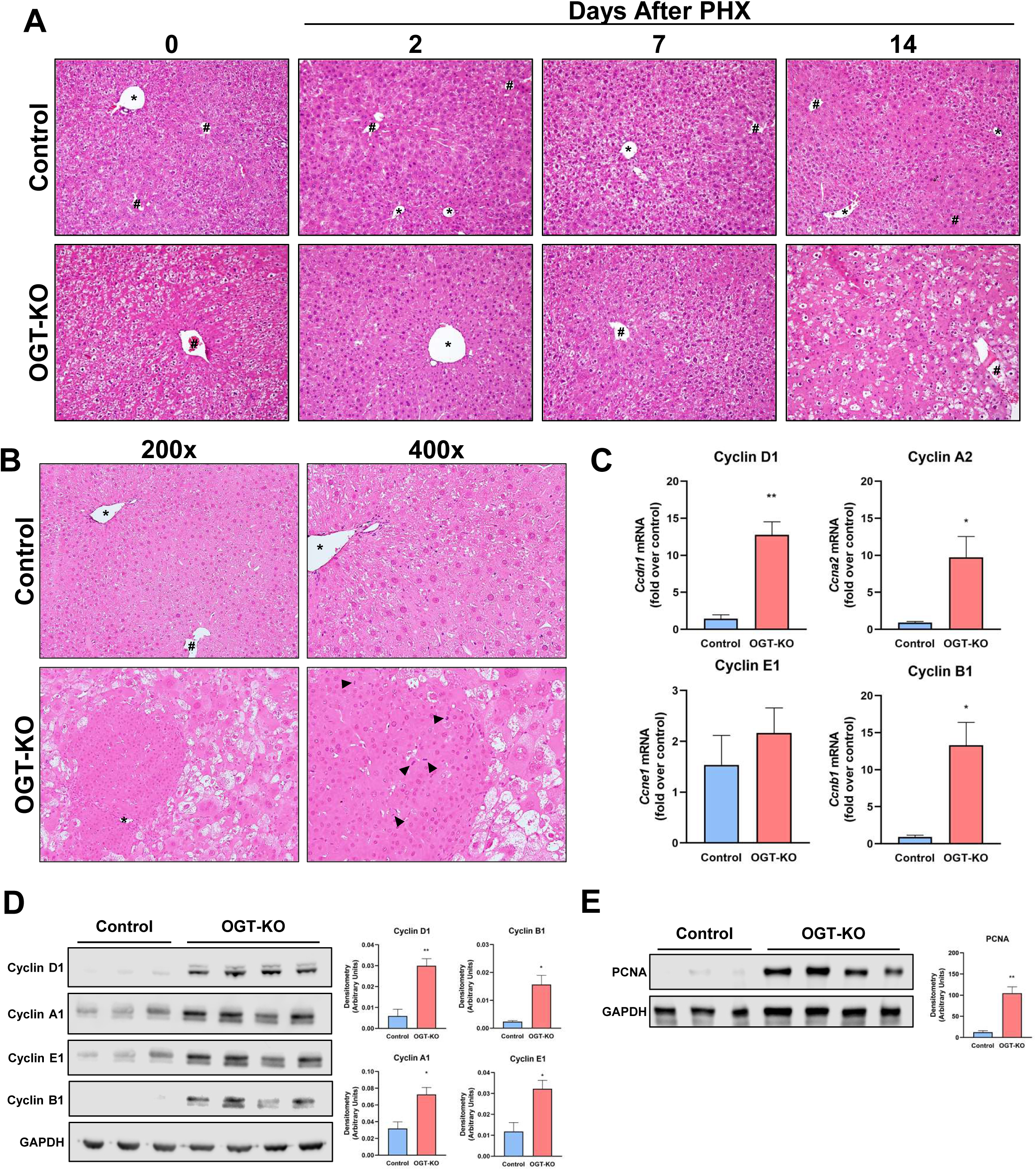
Hepatic Dysplasia is exhibited in OGT-KO mice 28 days after PHX. (A) Hematoxylin and eosin (H&E) representative photomicrographs of liver sections from control and OGT-KO mice 0 hour, 2, 7 and 14 days after PHX at 200x magnification. (B) H&E staining of 28 days after PHX at 200x and 400x magnification with arrow heads indicating mitotic figures. # represents central vein and * represents portal triad. (C) qPCR analysis of Cyclin D1, A2, E1 and B1 in OGT-KO and control livers from mice 28 days post-PHX, normalized to 18S and median of the control. Western blot analysis of liver lysate from 28 days after PHX OGT-KO mice of (D) cyclin D1, A1, E1, B1, and (E) PCNA with their corresponding densitometry. Bars represents the mean ± SEM, * p < 0.05 and ** p < 0.01.

Next, we investigated the fibroinflammatory changes in the livers of WT and OGT-KO 28 days after PHX. qPCR analysis revealed a significant induction in the expression of pro-inflammatory genes (*Tnfa, Il1b, Ifng* and *Il6*) and the anti-inflammatory gene *Il10*. OGT-KO mice showed induction in F4/80, a Kupffer cells marker (Fig. 4A), which was corroborated by immunofluorescence (IF) staining analysis showing increased F4/80^+^ cells in OGT-KO liver 28 days after PHX (Fig 5B). qPCR analysis indicated a significant increase in the hepatic stellate cell (HSC) marker desmin, its activation marker alpha-smooth muscle actin (*Acta2*) and *Tgfb*, a growth factor involved in activation of HSC (Fig. 4C). Consistent with these changes, OGT-KO livers also showed induction in fibrillar *Col1a1* and *Col1a2* expression (Fig. 4C). Picrosirius red (PSR) staining showed an increased collagen deposition in OGT-KO livers compared to the control (Fig. 4D). Moreover, hydroxyproline assay showed a significant increase in collagen deposition in OGT-KO livers 28 days after PHX (Fig. 4E). These parameters of inflammation and fibrosis did not show induction at any time point in OGA-KO mice (Fig. S4B-C). These data indicate that OGT-KO 28 days after PHX have significant inflammation and fibrosis.

**Figure 4.**
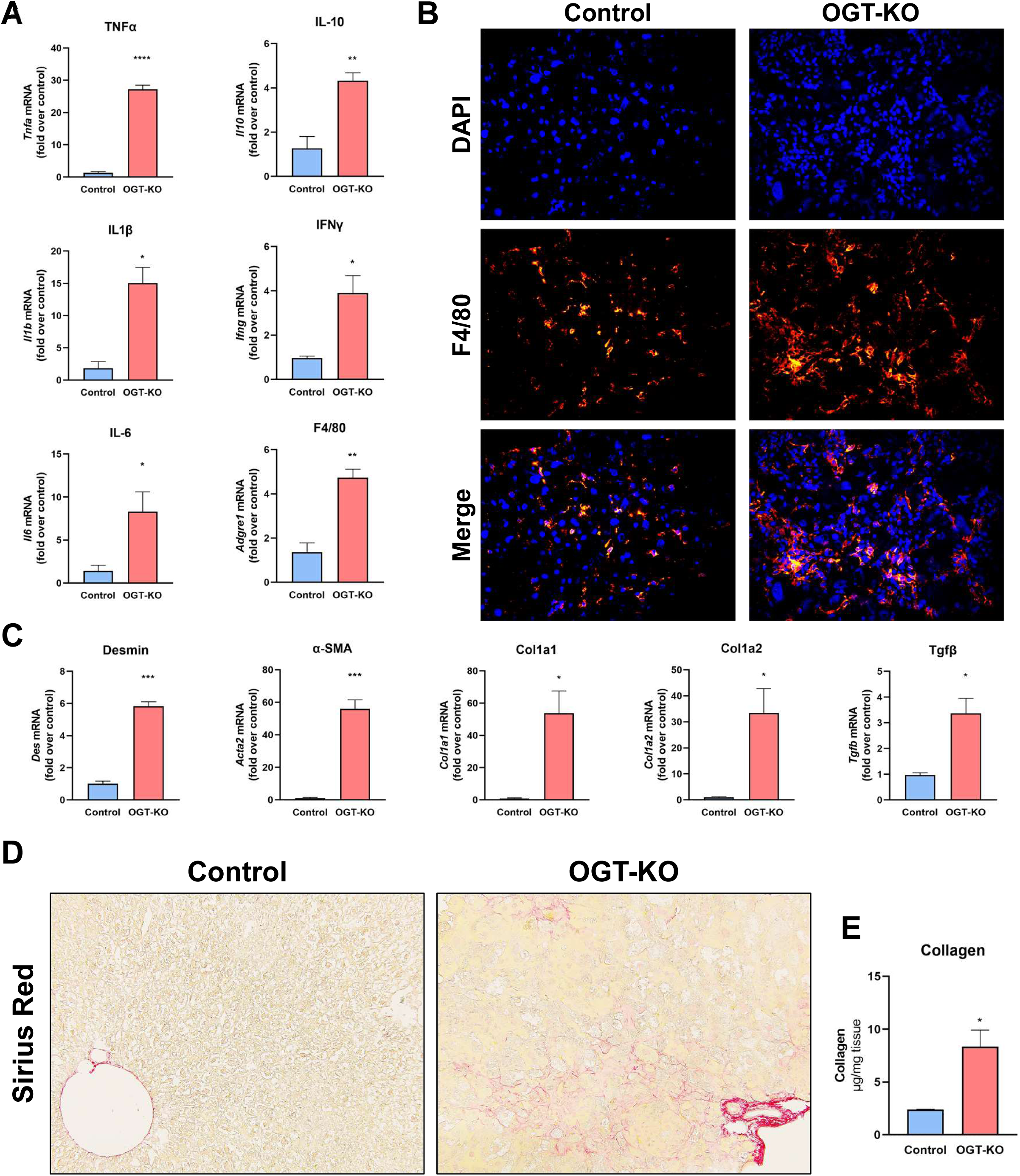
Increased inflammation and fibrosis in OGT-KO mice after completion of LR. (A) qPCR analysis of pro-inflammatory markers (*Tnfa, Il6, Il1b, Ifng*), anti-inflammatory marker (*Il10*) and Kupffer cell marker F4/80 (*Adgre1*). (B) Representative immunofluorescence images (400x) of F4/80 staining, DAPI and merged images of liver sections in OGT-KO mice 28 days after PHX. (C) qPCR analysis of markers of stellate cell activation *Des, Acta2, Tgfb, Col1a1* and *Col1a2* of livers from OGT-KO mice 28 days after PHX. (D) Photomicrographs (200x) of picrosirius red stained liver sections showing significant fibrosis (E) Hydroxyproline assay of liver tissue from OGT-KO mice 28 days after PHX. qPCR was normalized to 18S and the median of the control group. Bars represents the mean ± SEM, * p < 0.05, ** p < 0.01, *** p < 0.001 and **** p < 0.0001.

**Figure 5.**
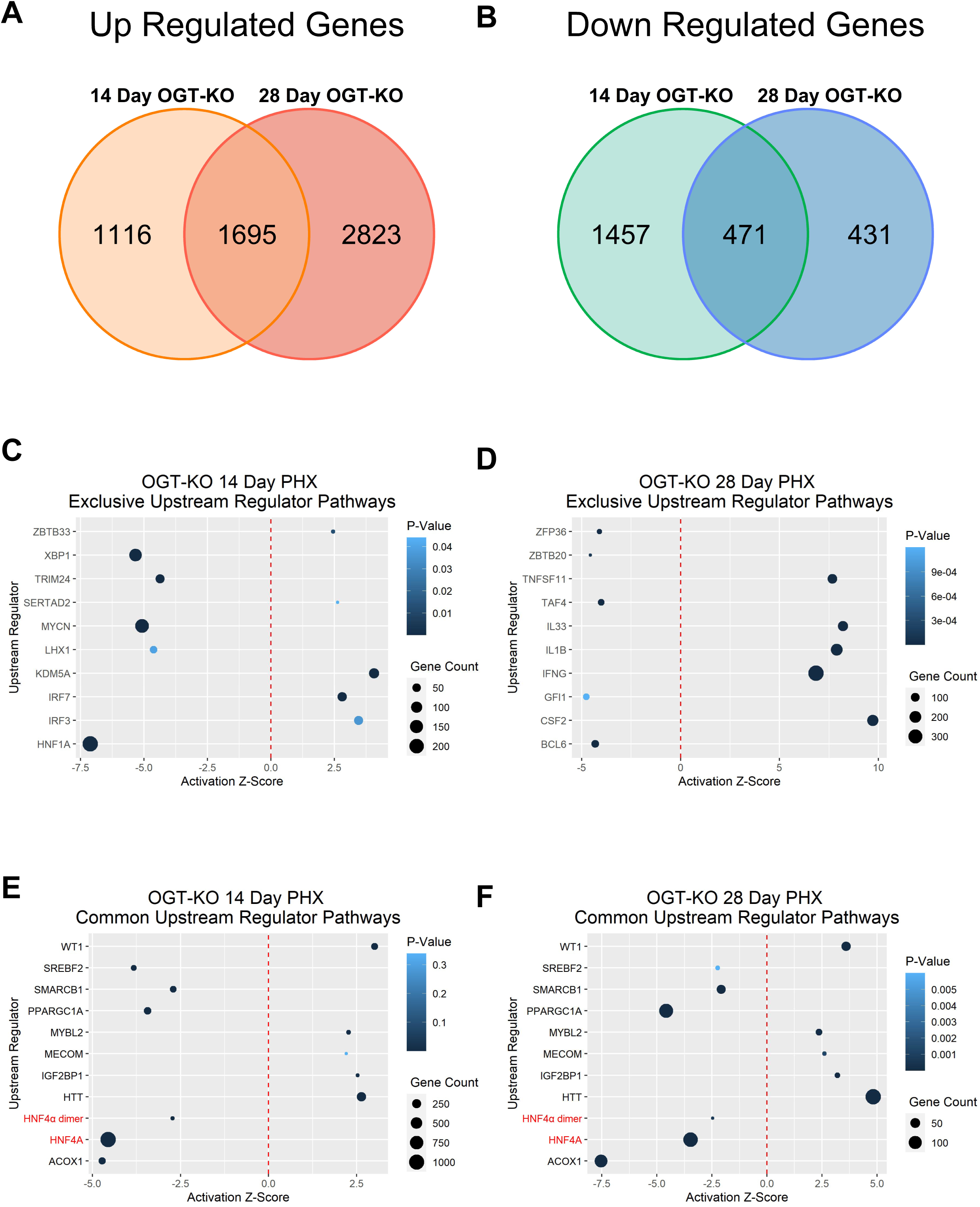
IPA of RNA-Seq data from 14 and 28 days after PHX revealed decrease activation of HNF4_α_. Venn diagrams showing (A) upregulated and (B) downregulated differentially expressed genes at 14 and 28 days after PHX using a 2-fold change cut off. Dot plots of upstream regulators that are common between (C) OGT-KO 14 days and (D) 28 days post PHX. The red dotted line indicates Z-Score of 0, the size of the dot represents the number of altered genes within that pathway and the color signifies the p-value.

### RNA-Seq Revealed Decrease Activation of HNF4_α_ in OGT-KO Mice 14 and 28 Days Post-PHX

To gain insight into the mechanisms of the defective termination of LR, we performed RNA-Seq on 14 and 28 days after PHX in OGT-KO and WT mice. Using a two-fold change expression cutoff, at 14 days after PHX, 2811 genes were upregulated and 1928 were downregulated. A total of 4518 genes were upregulated and 902 genes were downregulated at 28 days after PHX. 1,695 genes were commonly upregulated, and 471 genes were commonly downregulated between OGT-KO livers at 14- and 28-day after PHX (Fig. 5A-B, Table S1-2). Conversely, RNA-Seq studies on WT and OGA-KO mice at 14 and 28 days after PHX revealed only 19 commonly upregulated and 10 commonly downregulated genes (Fig. S4D, Table S3-4), which is consistent with no difference in liver regeneration between WT and OGA-KO mice.

An upstream regulator analysis was performed using Ingenuity Pathway Analysis (IPA) to identify key transcription regulators involved in the transcriptomic changes observed at 14 and 28 days after PHX. 14 days after PHX had exclusive activation of interferon regulatory factor 3 and 7 (IRF3, IRF7) and deactivation of HNF1A (Fig. 5C). Activation of IFNG, IL1B, and TNF were uniquely increased in 28 days after PHX, indicating an increase in inflammatory responses (Fig. 5D). Further, we identified transcription factors that were activated and inactivated at both 14 and 28 days after PHX in OGT-KO mice (Fig 6E-F). Interestingly, OGT-KO showed significant deactivation in HNF4_α_ function at both 14 and 28 days after PHX (Fig. 5E-F, Fig. S5A).

**Figure 6.**
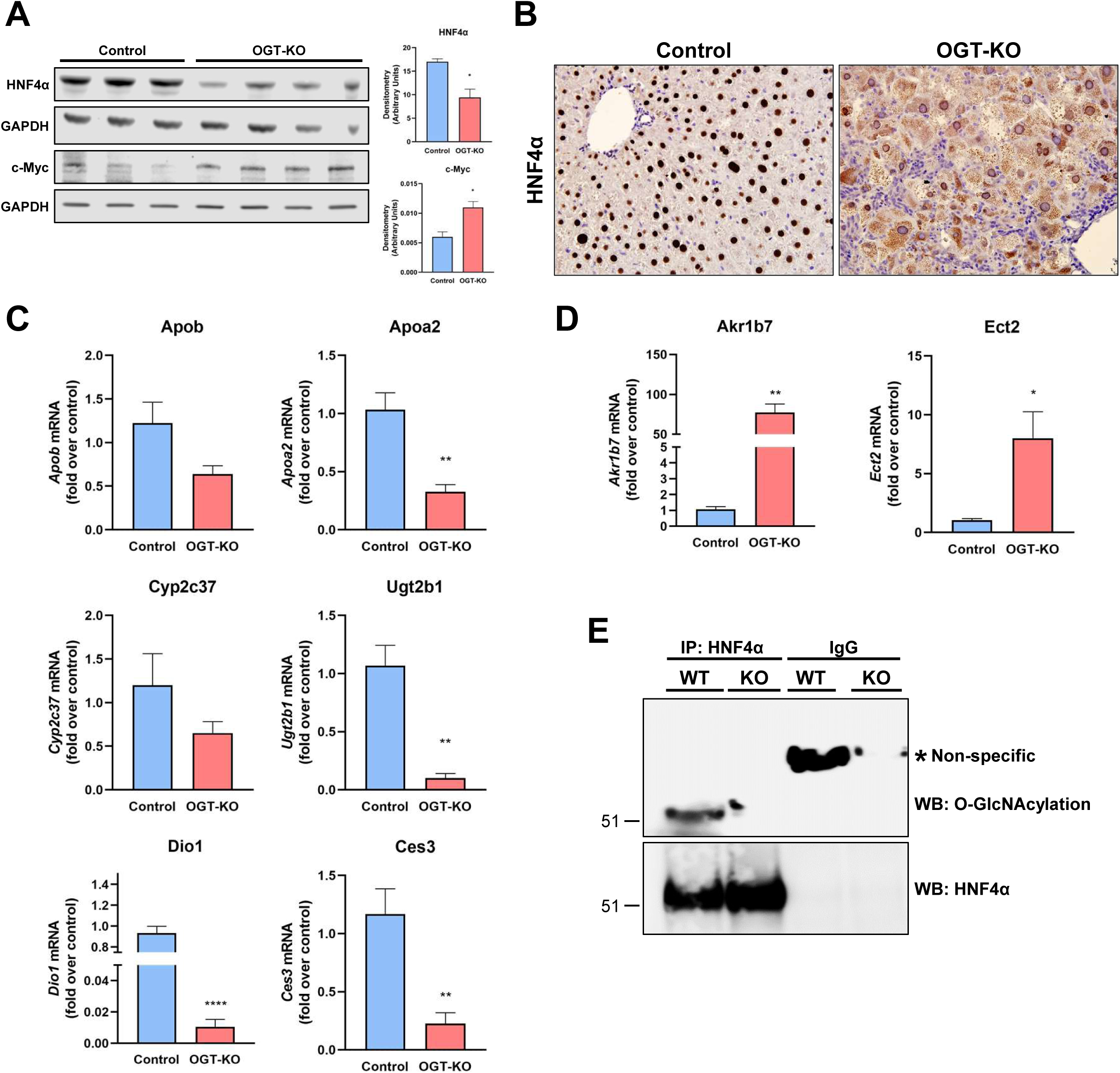
HNF4_α_ expression and function is decreased in OGT-KO mice 28 days after PHX. (A) Western blot analysis of HNF4_α_ protein levels in livers of control and OGT-KO mice 28 days after PHX, with corresponding densitometry. (B) Representative photomicrographs (400x) of HNF4_α_ IHC of control and OGT-KO mice 28 days post PHX. qPCR analysis of HNF4α (C) positive target genes and (D) HNF4α negative target genes at 28 days after PHX. qPCR was normalized to 18S and median of the control group. Bars represents the mean ± SEM, * p < 0.05, ** p < 0.01 and **** p < 0.0001. (E) IP pull-down of HNF4_α_ from WT and OGT-KO mice. Western blots analysis was performed on HNF4_α_ to show successful pulldown (upper blot) and O-GlcNAcylation (lower blot).

### HNF4α expression and activity declined in OGT-KO livers after PHX

Hepatocyte HNF4_α_ deficiency has been linked to spontaneous hepatocyte proliferation as well as defective termination of LR (15, 16). Therefore, we investigated HNF4_α_ expression and activity in OGT-KO mice 28 days after PHX. Western blot analysis of whole liver lysate indicated a significant decrease in HNF4_α_ protein levels in OGT-KO mice 28 days after PHX (Fig. 6A). Previous studies have shown that HNF4_α_ levels have reciprocal effects on the expression of the oncoprotein c-Myc (16, 17). Western blot analysis showed induction in c-MYC expression in OGT-KO livers at 28 days after PHX (Fig. 6A). Interestingly, IHC analysis revealed marked cytoplasmic redistribution of HNF4_α_ in OGT-KO mice (Fig. 6B). qPCR analysis of HNF4_α_ target genes indicated the loss of HNF4_α_ function with the decrease of positively regulated genes *(Apob, Apoa2, Cyp2c37, Ugt2b1, Dio1*, and *Ces3)* (Fig. 6C) and the increase of negatively regulated genes *(Akr1b7* and *Ect2)* (Fig. 6D). Finally, to determine if HNF4_α_ undergoes O-GlcNAcylation, immunoprecipitant (IP) experiments were performed. Total liver lysates from WT mice were used for immunoprecipitation of HNF4α protein and then Western blotting was conducted to detect the fraction of O-GlcNAcylated HNF4α. The data indicate that in WT normal livers, HNF4_α_ is heavily O-GlcNAcylated which is absent in OGT-KO mice (Fig. 6E).

### Methionine and Cysteine Metabolism is Altered in OGT-KO 14 and 28 Days After PHX

Lastly, we performed targeted metabolomics on WT and OGT-KO liver tissue lysates from 14 and 28 days after PHX. Principle component analysis (PCA) revealed that both WT and OGT-KO clustering was driven by both PC1 (31.7% variance explained) and PC2 (14.6% variance explained) and with each time point similar to each other (Fig. 7A). To determine altered metabolic pathways, Metabolomic Set Enrichment Analysis (MSEA) was performed on both significantly altered metabolites from 14- and 28-day samples. A significant impact on amino acid metabolism and glutathione metabolism was observed at 14 days after PHX in OGT-KO mice (Fig. 7B). Likewise, OGT-KO mice at 28 days after PHX had significantly altered energy substrate metabolisms such as pentose phosphate and glucose metabolism (Fig. 7C). Specifically, altered methionine synthesis was observed in OGT-KO mice at 28 days after PHX (Fig. 7C-E). The methionine synthesis metabolites, phosphorylcholine, choline, betaine, and methionine were significantly increased in the OGT-KO 28 days after PHX, whereas cysteine was significantly decreased (Fig. 7D-E). Importantly, HNF4_α_ regulates methionine and cysteine metabolism in hepatocytes (18).

**Figure 7.**
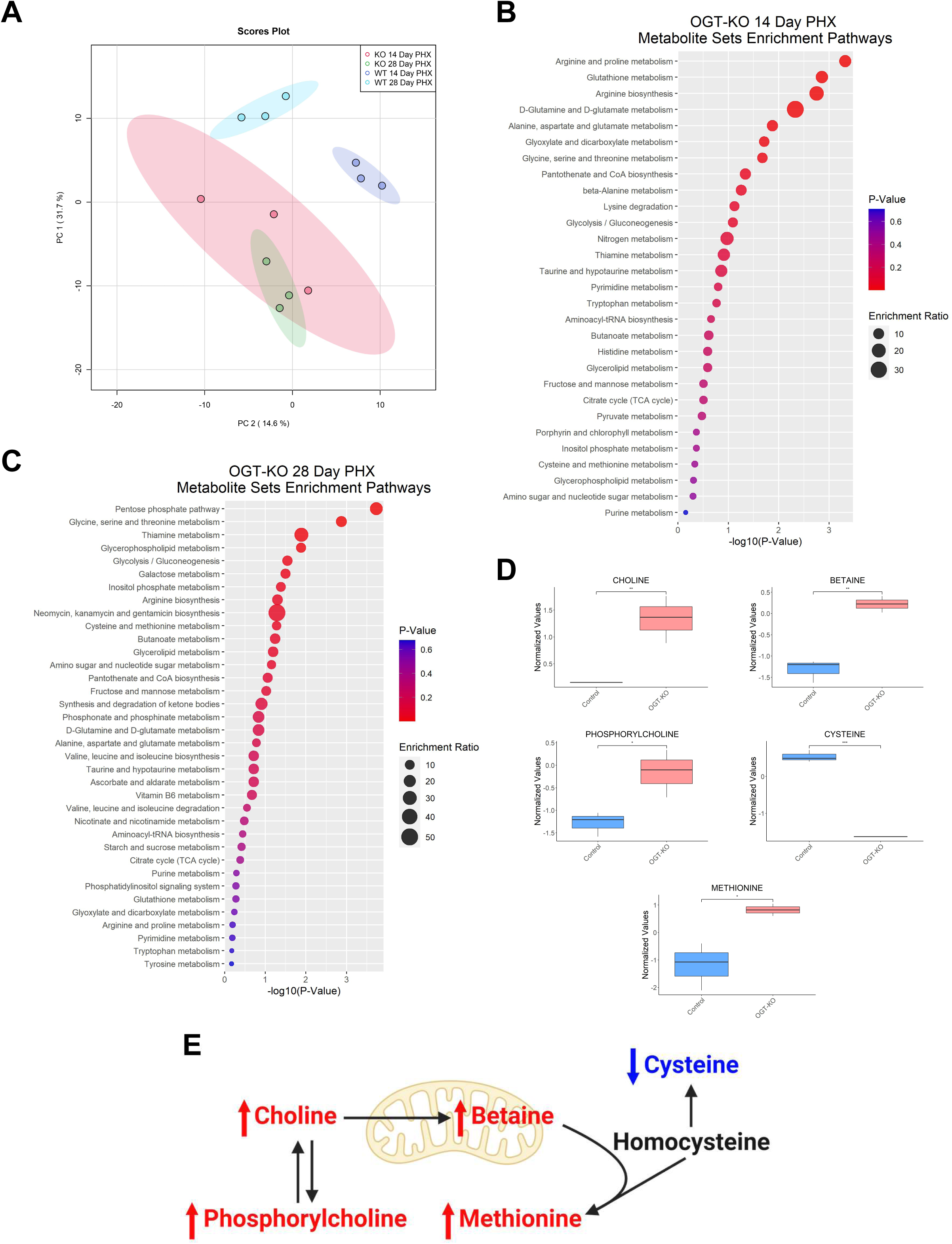
Metabolomic analysis show significant change in OGT-KO mice at 28 days after PHX. (A) PCA plot of control and OGT-KO mice for 14 and 28 days after PHX derived from targeted metabolomics. Ellipses represent clusters for each group. Metabolite Set Enrichment Analysis of OGT-KO mice (B) 14 days and (C) 28 days after PHX, color represents p-value and size corresponds to enrichment ration. (D) Significantly altered metabolites in methionine and cysteine metabolism in OGT-KO mice 28 days after PHX. Bars represents the mean ± minimum and maximum normalized value, * p < 0.05, ** p < 0.01 and **** p < 0.0001. (E) Metabolic pathway of methionine and cysteine with red and blue indicated increased or decreased metabolite in OGT-KO mice 28 days after PHX, respectively.

## Discussion

Liver regeneration (LR) after PHX, in mice, can be divided into three broad phases. The initiation of regeneration is where hepatocytes and other cells partially dedifferentiate and enter cell cycle, progression of regeneration is where cells go through mitosis and divide, and finally, the termination of regeneration is when cell proliferation declines, newly divided cells re-differentiate, and liver tissue is reorganized. The initiation and progression phases are relatively rapid and completed within 5 days after PHX. Termination of regeneration is more prolonged and can take up to a week. Most of the liver mass and function is restored by 2 weeks (19, 20). After PHX, multiple signals have been well characterized that initiate the cell cycle in LR, termed primary mitogens, such as the hepatocyte growth factor (HGF) and epidermal growth factor (EGF) (19, 21-23). Auxiliary mitogens, consisting of TNF_α_ and IL-6, enhance the prolific effects of the primary mitogens during the initiation and progression of LR (19, 24). However, much less is known about the termination of LR. Integrin-linked kinase (ILK), critical for hepatocyte interactions with extracellular matrix (ECM), has been identified as a termination signal (14, 25). Recent studies from our laboratory have shown that HNF4_α_, the master regulator of hepatic differentiation (26), is critical for re-differentiation of hepatocytes after proliferation and plays a critical role in termination of liver regeneration (16, 17, 27). Previous studies have shown that defective termination of LR will result in significant hepatomegaly and death due to decline in liver function. We hypothesize that in cases where animals do not die due to acute liver failure, defective termination of liver regeneration will lead to hepatic dysplasia that can progress to liver cancer. Evidence presented here supports this hypothesis and indicates that hepatic O-GlcNAcylation is a major regulator of the termination of liver regeneration. After PHX, OGT-KO mice exhibited no differences during the initiation phase and progression of LR. However, during the termination of LR, we found sustained proliferation in hepatocytes indicating a termination defect (Fig. 2A). This was further exhibited by the significant increase in liver-weight to body-weight ratio 14 and 28 days after PHX (Fig. 1D). The foremost change was the development of hepatic nodules accompanied by significant inflammation and early fibrotic changes.

These data are consistent with previous observations that O-GlcNAcylation plays a critical role in cell cycle regulation (8, 10, 28, 29). O-GlcNAcylation of proteins increases during the progression of G_1_ phase following a rapid decrease when cells enter S phase (9). This is contributed to an immediate induction of OGA protein levels during S phase (9). Throughout G_1_ phase, Rb is O-GlcNAcylated and when the G_1_/S-checkpoint into S phase occurs, Rb is needed to be phosphorylated indicating cross-talk of PTMs during cell cycle progression (30). After S phase, during the G_1_ to M phase transition, increasing O-GlcNAcylation by OGA inhibition delays the advancement to M phase (10). During mitosis, global O-GlcNAcylation levels decrease (10). Either increasing or decreasing O-GlcNAc during mitosis will lead to aberrant spindle formation causing mitotic defects (8, 31). We found that OGT deletion (reduction of O-GlcNAcylation) in hepatocytes for seven days resulted in mild hepatomegaly due to hyperplasia as well as hypertrophy (Fig. 1D, B). Importantly, this indicates that OGT regulates hepatocyte proliferation without induction of proliferation by PHX.

We found ballooning hepatocytes accompanied with significant liver injury in OGT-KO mice at 14 and 28 days after PHX (Fig. 1C). Also, initial liver injury at 2 days post-PHX was higher in OGT-KO mice as compared to WT. These data indicate that loss of O-GlcNAcylation may make hepatocytes more vulnerable to cell death. These data are consistent with previous studies by Zhang et al demonstrated that OGT deletion, in the liver using a CRE governed by the albumin promoter, resulted in significant necroptosis and fibrosis (7). They further showed that RIP3K undergoes O-GlcNAcylation which reduces the stability of the protein (7). Our studies showed that delayed cell death in OGT-KO mice after PHX is both necroptosis as well as apoptosis (Fig. S2A-C). Interestingly, we observed an increase in p62, a marker of autophagy and Mallory-Denk bodies, in OGT-KO mice after PHX, which is commonly found in metabolic disorders and hepatocellular neoplasms (Fig. S2E) (32). It is well known that autophagy inhibits apoptosis which suggests that the induction of p62 is probably due to Mallory-Denk body formation (33). To determine if autophagy is contributing to cell death, more studies will need to be done. However, the exact mechanisms of increased spontaneous cell death at two weeks after PHX in OGT-KO mice are not known. It is plausible that lack of hepatic redifferentiation, which is a critical component of termination of regeneration, may trigger the cell death at such a delayed time point.

This argument is supported by the RNA-seq studies on 14- and 28-day time points, which revealed significant decline in HNF4α target gene expression (Fig. S5A). HNF4_α_ is extremely critical in maintaining hepatocyte function, and we have previously shown that deletion of HNF4α results in defective termination of liver regeneration (16, 27). HNF4α protein levels and its function are regulated by post-translational modifications. Most PTMs of HNF4_α_ reduce function and stability. For instance, the C-terminal of HNF4_α_ undergoes SUMOylation to promote degradation (34), acetylation of K458 attenuates the transcriptional activity, deactivating HNF4_α_ (35) and phosphorylation occurs on multiple residues. Other studies have shown that protein kinase A (PKA)-mediated S142 phosphorylation decrease HNF4_α_ protein levels (36). Extracellular signal⍰regulated protein kinase 1/2 (ERK1/2) has been shown to phosphorylate multiple residues on HNF4α including S138/T139, S143, S147/S148, S151, T166/S167, and S313 (37). Interestingly, the residue S142 was further found to be phosphorylated by ERK1/2 indicating two kinases acting on the same residue (37). Here, we show that HNF4_α_ is O-GlcNAcylated and without this, HNF4_α_ function is lost. It is known that serine or threonine residue can be alternatively phosphorylated or O-GlcNAcylated (38). Our studies give rise to the possibility that normally HNF4α is O-GlcNAcylated, which prevents its degradation or cytoplasmic relocalization by inhibiting its phosphorylation. If this dynamic relationship occurs, this could explain why increasing O-GlcNAcylation in OGA-KO livers had no physiological phenotype. Further studies will need to be done to map the specific O-GlcNAcylated residue(s) on HNF4α. Nonetheless, our studies have uncovered a novel mechanism by which HNF4α stability and function are regulated. Further, our studies show that O-GlcNAc mediated regulation of HNF4α function is critical in termination of liver regeneration.

When OGA or OGT is deleted, a compensatory effect occurs, altering the expression of the reciprocal enzyme (39). We observed this phenomenon for the first time in hepatocyte-specific OGT-KO and OGA-KO mice (Fig. 1A-B). In OGT-KO mice, this compensatory effect cannot increase O-GlcNAcylation levels. However, in OGA-KO, the compensatory effect of decreasing OGT levels will reduce global O-GlcNAcylation closer to basal levels. This could potentially mitigate the adverse effects of exorbitant O-GlcNAcylation and explain why OGA-KO mice had normal LR. Future experiments will need to be done to artificially increase O-GlcNAcylation to prevent the compensatory effects in OGA-KO mice to determine the effects of augmented O-GlcNAcylation.

In summary, our studies are the first to examine and manipulate O-GlcNAcylation over a time course throughout the LR process after PHX. Our findings suggest that lack of O-GlcNAcylation causes defects in the termination of liver regeneration. Ablation of O-GlcNAcylation impedes the function and stability of HNF4_α_, which leads to decreased hepatocyte redifferentiation, significant necroinflammation, early fibrotic changes and formation of dysplastic nodules. These results confirm the role of O-GlcNAcylation in termination of regeneration and preventing hepatic dysplasia and highlight O-GlcNAcylation as a therapeutic target in hepatic regenerative medicine.

## Grant support and Acknowledgments

These studies were supported by NIH-COBRE (P20 RR021940-03, P30 GM118247), NIEHS Toxicology Training Grant (T32 ES007079-34), NIH R01 DK0198414 and NIH R56 DK112768

## Notes

### Competing Interest Statement

The authors have declared no competing interest.

